# Mechanisms governing irritant-evoked activation and calcium modulation of TRPA1

**DOI:** 10.1101/2019.12.26.888982

**Authors:** Jianhua Zhao, John V. Lin King, Candice E. Paulsen, Yifan Cheng, David Julius

## Abstract

The TRPA1 ion channel is a chemosensory receptor that is critical for detecting noxious chemical agents that elicit or exacerbate pain or itch. Here we use structural and electrophysiological methods to elucidate how a broad class of reactive electrophilic irritants activate TRPA1 through a two-step cysteine modification mechanism that promotes local conformational changes leading to widening of the selectivity filter to enhance calcium permeability and opening of a cytoplasmic gate. We also identify a calcium binding pocket that is remarkably conserved across TRP channel subtypes and accounts for all aspects of calcium-dependent TRPA1 regulation, including potentiation, desensitization, and activation by metabotropic receptors. These findings provide a structural basis for understanding how endogenous or exogenous chemical agents activate a broad-spectrum irritant receptor directly or indirectly through a cytoplasmic second messenger.

## Introduction

The ability to recognize noxious chemicals constitutes a key aspect of pain sensation, one that is critical for avoiding environmental toxicants or detecting endogenous products of tissue injury. The TRPA1 ion channel (also known as the ‘wasabi receptor’) is expressed by primary afferent nerve fibers, where it functions as a low threshold sensor for reactive electrophiles of widely disparate chemical structures ranging from small volatile environmental irritants like acrolein or allicin to endogenous reactive lipids, such as 4-hydroxy-2-nonenal and 15-deoxy-Δ(12,14)-prostaglandin J(2)^1-6^. In contrast to the classic lock-and-key model of receptor-ligand interaction, electrophiles activate TRPA1 through an unusual mechanism involving covalent modification of specific nucleophilic residues within the channel’s cytoplasmic amino terminus^7,8^. TRPA1 is also a ‘receptor-operated’ channel that is activated downstream of metabotropic receptors that couple to phospholipase C (PLC) signaling pathways, most notably by release of calcium from intracellular stores^9,10^.

In our initial structural analysis of TRPA1, we identified an intricately folded cytoplasmic region (dubbed the allosteric nexus) situated just below the transmembrane core of the channel that contains three cysteine residues implicated as functionally relevant sites for electrophile modification^11^. However, we were not able to visualize such modifications or capture the channel in distinct conformational states, limiting our understanding of how TRPA1 recognizes and responds to an incredibly broad spectrum of reactive irritants. Nor did we gain insight into structural mechanisms underlying calcium-dependent channel regulation, an important functional attribute of TRPA1 and other members of the extended TRP channel family. In this study, we capture TRPA1 in apo and liganded states, addressing these and other questions that are key to understanding channel behavior and mechanisms of drug action.

## Results

### Dynamic equilibrium between two states

Like many TRP channels, TRPA1 exhibits ‘flickery’ behavior characterized by spontaneous openings or rapid transitions between open and closed states in the presence of agonists (Fig. 1A, and fig. S1A)^12,13^. We used single particle cryo-EM to capture these distinct states within a single sample in the absence or presence of ligands, thereby identifying two main conformations that could be assigned to specific functional states based on these pharmacological criteria. Thus, one main state predominated in the presence of iodoacetamide (IA), an electrophilic agonist that forms an irreversible covalent adduct to the channel, and we therefore designated this as an activated state (Fig. 1B; and fig. S2 and S3)^7,8^. The other state bound the antagonist, A-967079 (A-96) ^14^, and was designated as closed. Whereas IA was added to TRPA1-expressing cells prior to channel purification, A-96 was added after detergent extraction, likely accounting for the fact that the antagonist specifically recognized but did not shift distribution of particles towards the closed state (Fig. 1A).

**Figure 1.**
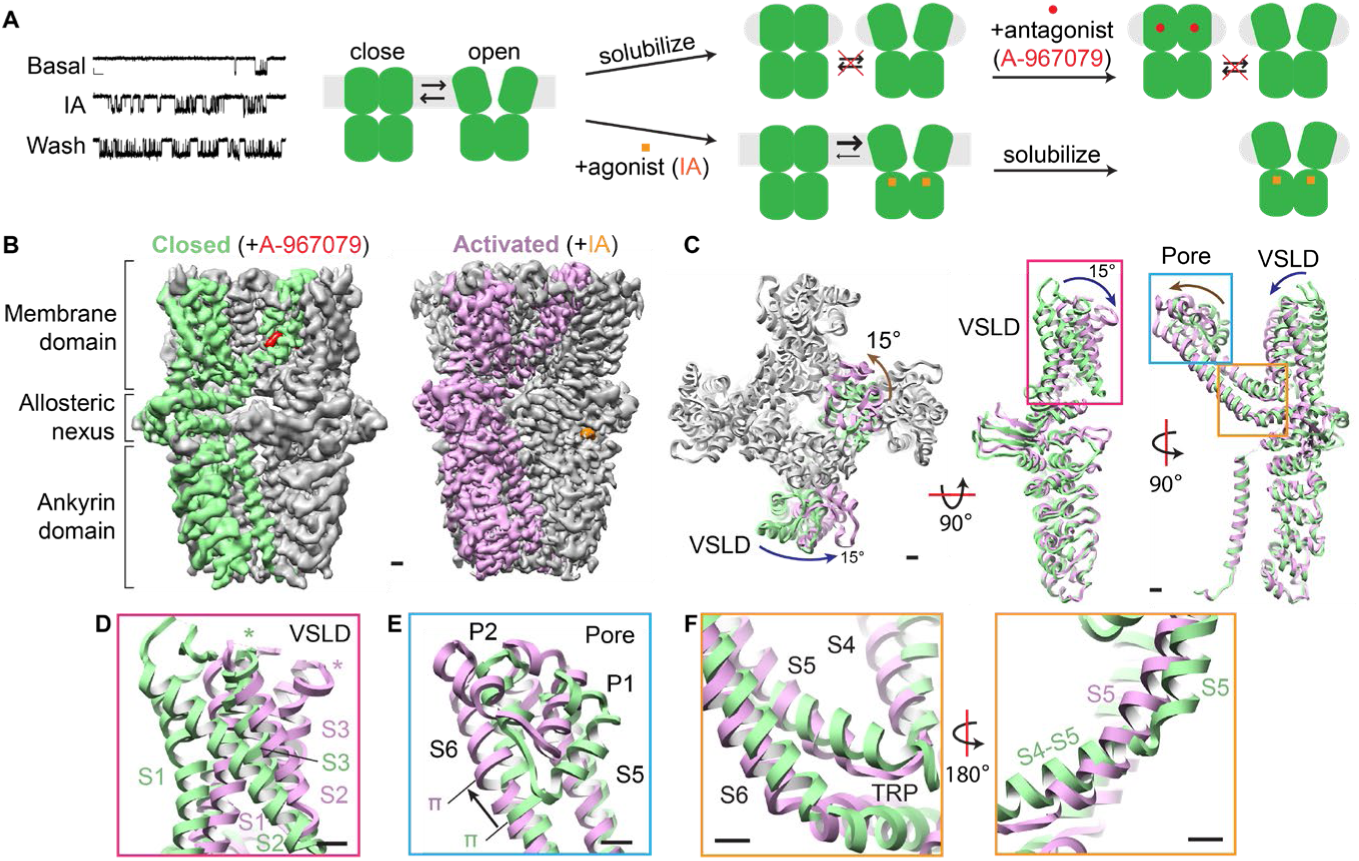
Dynamic equilibrium between closed and activated conformations. **(A)** Closed and open states of TRPA1 were captured by incubation with an irreversible electrophilic agonist (Iodoacetamide, IA) or an antagonist (A-967079, A-96) before or after membrane solubilization, respectively. Inset: representative single channel traces showing spontaneous (basal) and IA (100 µM)-evoked (persistent) TRPA1 channel openings (V_hold_ = −40mV; scale bars: x = 10msec, y = 2pA. **(B)** Cryo-EM density maps of TRPA1 bound to A-96 or IA in the closed or activated state, respectively. A-96 binds to the membrane domain while iodoacetamide binds to the allosteric nexus. **(C)** Comparison between subunits in closed and activated conformations, showing ∼15° rotation of the voltage sensor-like domain and twisting and translation of the pore domain. The ankyrin repeat domain remains stationary between the two conformations. **(D)** The VSLD rotates around the cytoplasmic base of transmembrane α-helices S1 and S2 in a near-rigid-body movement. **(E)** The pore domain twists and translates upward towards the extracellular milieu, concomitant with a shift in the π-helix of S6 by one helical turn. **(F)** Conformational changes in the upper pore region are coupled to widening of the lower gate through straightening of the S5 α-helix, enabling movement of S6 to facilitate gating. Scale bars: 10Å.

Comparison of the structures of these two states revealed significant conformational differences within the transmembrane core and the membrane-proximal allosteric nexus (Fig. 1C – F). The entire transmembrane region rotates by ∼15° relative to the stationary cytoplasmic ankyrin repeat domain (Fig. 1C). Specifically, the voltage sensor-like domain undergoes a near ridged body rotation (Fig. 1D), which is accompanied by rotation and upward translation of the pore loop and pore helices (Fig. 1E). These transitions are linked by movements in the S5 and S6 helices: the π-helix in S6 shifts up by one helical turn and, in concert with S5, causes an upward shift of pore helices P1 and P2 (Fig. 1E); the S4-S5 linker and S5 transmembrane α-helix straighten into a single α-helix (Fig. 1F), which coordinates movement of the pore helices and S6 to couple the upper and lower gates. Taken together, these transitions are reminiscent of those between closed and desensitised states observed in the cool/menthol receptor TRPM8, but dissimilar to the more local conformational changes that accompany gating in the heat/capsaicin receptor TRPV1^15-18^.

### Coupling between upper and lower gates

Profiles of the ion permeation pathway showed a substantial difference in restriction sites at the levels of both the upper gate/ion selectivity filter and lower gate (Fig. 2A and B). We see a marked widening of the lower gate formed by residues Ile957 and Val961, dilating the pore from 5.3 to 7.8 Å in diameter (Fig. 2C) and increasing its hydrated radius from 0.9 to 2.1 Å (Fig. 2B). These changes are nearly identical of those seen in TRPV1 when comparing apo and capsaicin-activated states, indicative of transition from a fully closed to an open state and expected for a true gate that controls ion flux^16-18^. Local resolution analysis of the activated TRPA1 map shows that the lower gate has slightly lower resolution compared to surrounding regions (fig. S3), suggesting that this region is more dynamic in the activated versus closed state.

**Fig. 2.**
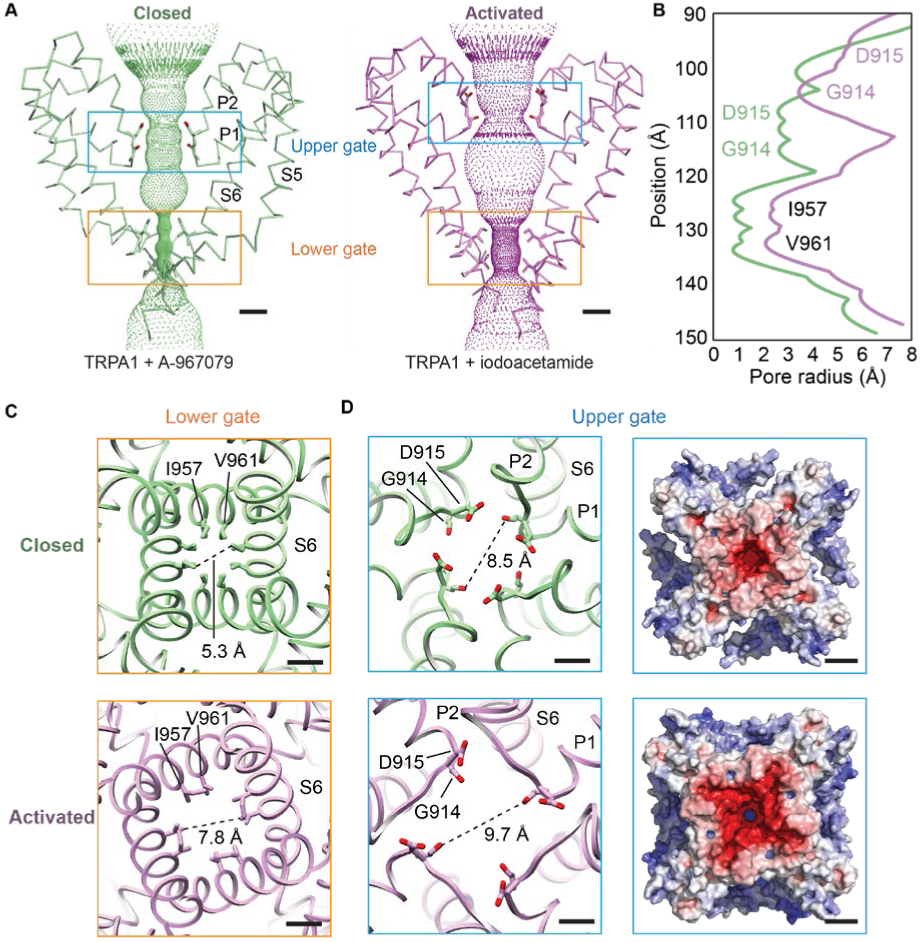
Coupled dilation of upper pore region and lower gate. **(A)** Activation of TRPA1 involves concerted dilation of both the upper and lower gates along with an upward shift of the selectivity filter. **(B)** The lower gate is formed by residues Ile957 and Val961 while the upper gate is formed by Asp915 and the backbone carbonyl of Gly914. **(C)** Widening of the pore is associated with counter-clockwise rotation of the transmembrane α-helices (viewed from the extracellular face to the cytoplasmic side). **(D)** Acidic residues lining the upper pore region create a highly negatively-charged surface, especially in the activated conformation of TRPA1, likely facilitating selectivity for calcium. Scale bars: 5 Å. Electrostatic surface charge distribution of TRPA1’s cytoplasmic face in the apo- and activated states were calculated in APBS^47^ and are displayed at ±10 kT/e^-^. Scale bars: 25Å.

At the upper restriction site, movement of pore helices P1 and P2 results in an upward shift and dilation of the upper gate/selectivity filter formed by residues Gly914 and Asp915 (Fig. 2A and B). This alters the overall profile of the outer pore wall from a V- to a U-shaped funnel, which is accompanied by an increase in the restriction diameter from 8.5 to 9.7Å (Fig. 2D). Remarkably, widening of both upper and lower pore restrictions is evoked by attachment of an electrophile to the allosteric nexus, which sits below the transmembrane core, arguing for an allosteric coupling mechanism between this nexus and the two gates. Regarding communication between the gates, we observe that S5 in the closed state contains a small bend that straightens upon transition to the open state (Fig. 1F), consequently lifting and rotating the pore helices backward to alter configuration of the selectivity filter (Fig. 2A). Consistent with these proposed coupling movements, the A96 antagonist wedges into a pocket close to the S5 bend and just beneath the pore helices (fig. S1E and S9), as previously described^11^. We can now infer the mechanism of A96 action: sitting at the elbow of S5, this inhibitor likely prevents opening of the lower gate by locking S5 in its closed (bent) conformation, thereby preventing gating movements of the S5-S6 pore module (Fig. 5C and fig. S9).

What might be the physiologic relevance of such changes at the level of the outer pore region, where substantial shape changes occur, but with a less dramatic alteration in restriction diameter than observed in TRPV1? Interestingly, this transition involves upward and outward rotation of Asp 915 (Fig. 2B and D), a residue that has been previously shown to influence calcium selectivity of activated TRPA1 channels^19,20^. The net result of this rotation is to increase the negative electrostatic surface potential in the outer pore region (Fig. 2D), consistent with enhanced calcium preference in the activated state^19,21-23^. Indeed, Asp915Ala or Asn mutants do not show a preference for calcium^19^ and models of these mutants show reduced negative electrostatic surface potential in the activated state (fig. S4). Taken together, these observations support the notion that conformational transitions at the level of the selectivity filter account for dynamic ion selectivity of TRPA1 channels^19,21-23^.

### Two-step mechanism of electrophile action

Our activated TRPA1 structure was obtained using IA, which functions as an irreversible agonist by forming stable covalent adducts with cysteines (Fig. 1A, fig. S1A and S6A)^7,8^. We observed clear attachment of IA at just one position, Cys621 (fig. S5A), consistent with the exceptional nucleophilic character of this residue^11,24^. Strikingly, this modification also stabilized an ‘activation loop’ (which we now dub the A-loop) in an upward configuration, exposing a pocket containing the modified residue. To better visualize agonist attachment and conformational rearrangements of the A-loop, we visualized the channel following modification with a bulky version of IA (BODIPY-IA, BIA) (fig. S1E). Furthermore, we found that solubilization with the detergent LMNG resulted in overall higher resolution while still permitting conformational changes within the allosteric nexus. Consequently, we visualized the apo channel with an improved overall resolution of ∼3.1 Å (fig. S2 and table S1), enabling us to see the A-loop in a clearly defined downward configuration, largely occluding the pocket containing Cys621 in the unmodified form (Fig. 3A and fig. S3). The BIA-labeled structure was also visualized with substantially improved resolution (2.6 Å) (fig. S2), where the larger density of BIA confirmed Cys621 as the primary nucleophilic modification site (Fig. 3B and fig. S5B). While other cysteines within this region (most notably Cys665) contribute to channel activation, they are modified at substantially reduced rates compared to Cys621^24^. Indeed, in the presence of IA we observed a weaker density associated with Cys665, which may reflect partial modification at this site (fig. S5a).

**Fig. 3.**
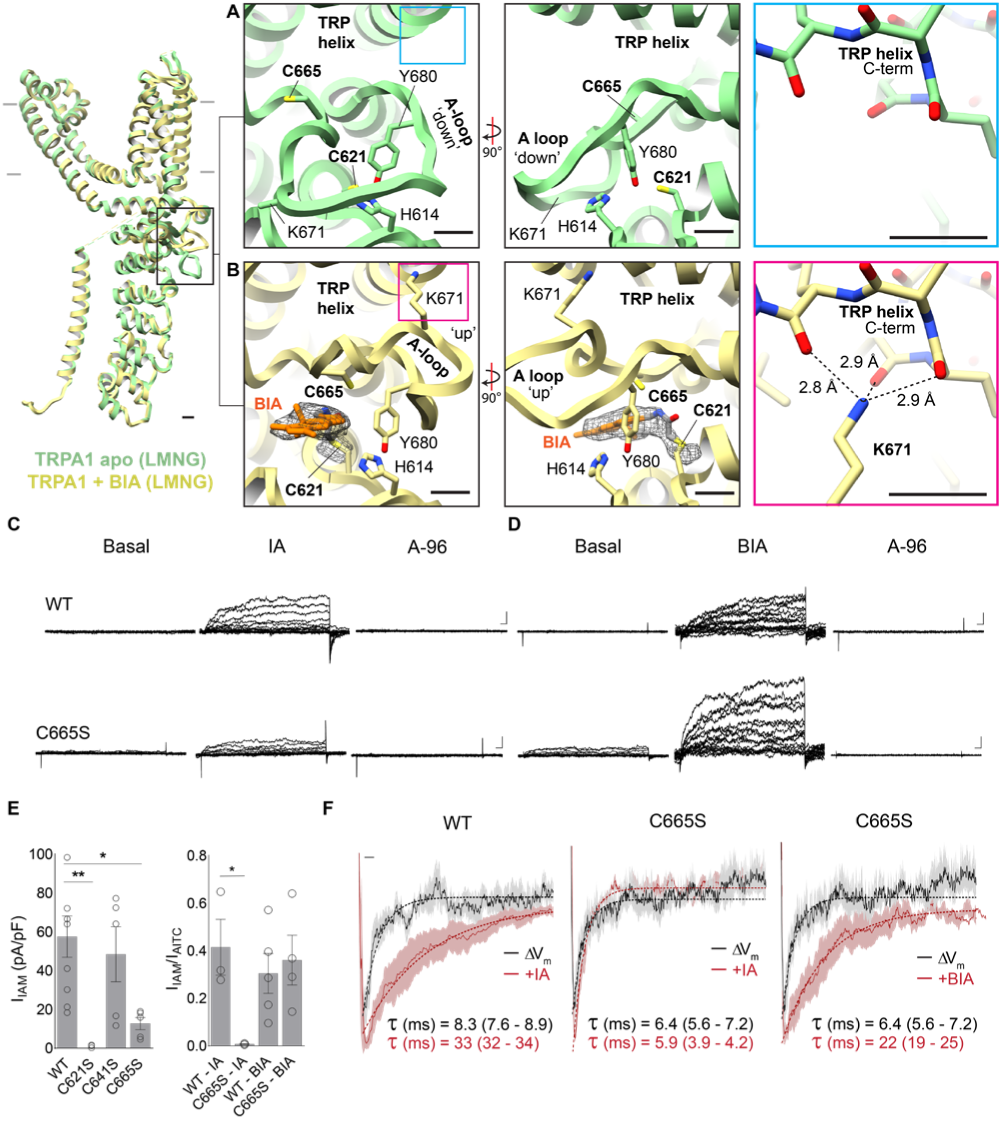
Activation by electrophiles occurs through a two-step mechanism. **(A)** A dynamic activation loop (A-loop) adopts a ‘down’ conformation in the apo channel, partially occluding a reactive pocket containing Cys621. **(B)** Following attachment of BODIPY-IA (BIA) to Cys621, the A-loop transitions to an ‘up’ conformation, bringing Cys665 into the reactive pocket and repositioning Lys671 to coordinate backbone carbonyl oxygens at the TRP domain C-terminus. Scale bars: 5 Å. **(C)** IA (100 μM) or **(D)** BIA (100 μM)-evoked whole-cell currents for WT and Cys665Ser mutant TRPA1 channels (500msec. voltage steps from −80 to 80 mV; n = 5-9 independent experiments/construct; scale bars: x = 25msec., y = 100pA). **(E)** (left) Quantification of IA-evoked WT and mutant TRPA1 currents at 80mV (n = 5-9 independent experiments/construct; *, *p* < .05; ** *p* < 0.01, one-way ANOVA with post-hoc Holm-Sidak correction for multiple comparisons (left panel). (right) Comparison of BIA (100 μM) to AITC (250 μM)-evoked currents for TRPA1 WT and Cys665 Ser mutant channels (n = 3-5 independent experiments/construct/treatment; *, *p* < 0.05, one-way ANOVA with post-hoc Holm-Sidak correction for multiple comparisons. **(F)** Scaled averaged IA (100 μM) or BIA (100 μM)-evoked tail currents for TRPA1 WT and Cys665Ser mutant channels. Mean deactivation time constants (τ) are shown with 95% CI in parentheses; n = 4-6 independent experiments per construct. Pre-pulse: 80mV, 500msec. Test pulse: - 120mV, 250msec. Scale bar: x = 5msec., y = arbitrary units.

Interestingly, Cys665 rotates into the pocket containing modified Cys621 as the A-loop is stabilized in the upward configuration (Fig. 3A and B). This reorientation of Cys665 is predicted to lower its pK_a_ (from 11.2 to 8.8) and enhance its nucleophilicity (fig. S6J) suggesting that modification at this site also occurs in the process of channel activation. The necessity for modification at one or both cysteines may depend on electrophile properties and whether the size or charge of the modification is sufficient to alter configuration of the reactive pocket. To test this idea, we mutated three cysteines (Cys621, Cys641, and Cys665) within the vicinity of the reactive pocket and used patch clamp recording to assess their sensitivity to IA and its bulkier cousin, BODIPY-IA. All double mutant combinations were IA insensitive (fig. S6B-D), showing that Cys621 alone is insufficient to support channel activation by this small electrophile (even though Cys621 was fully labeled by BODIPY-IA in Cys 41 Ser, Cys665 Ser double mutants, fig. S6I). We next examined single cysteine substitutions and found that Cys641 Ser behaved similarly to wild type channels (Fig 3E, fig. 6F), whereas Cys621 Ser was IA insensitive (Fig. 3E, fig. S6E) and Cys665Ser retained ∼30% sensitivity (Fig. 3C and E) This is consistent with a two-step model whereby Cys621 is a primary site of electrophile modification, priming A-loop reorientation and modification of Cys665 to elicit full channel activation.

Further evidence for this model comes from measurements of channel closure times, which reflects the coupling efficiency between agonist binding and channel gating. To assess this parameter, we examined tail currents following activation of channels by membrane depolarization in the absence or presence of irreversible agonists. For wild type TRPA1 channels, the rate of channel closure was slowed ∼3-fold by IA (Fig. 3F and fig. S6G) (*p* < 0.05, ratio paired two-tailed student’s *t*-test), as expected if electrophile modification enhances coupling efficiency of A-loop reorientation to channel gating. For Cys665Ser mutants, which retain appreciable sensitivity to IA, the rate of channel closure was similar to that determined in the absence of agonist (Fig. 3F and fig. S6G), suggesting that modification at Cys665 is important for stabilizing and coupling A-loop reorientation to gating. In the case of BODIPY-IA, however, we see attachment to only one site, suggesting that bulky electrophiles stabilize this active loop conformation by modifying Cys 621 alone. Remarkably, the single Cys665Ser mutant showed full sensitivity to BIA with regard to both response magnitude and tail current decay time (Fig. 3D, E, and F; and fig. S6H) (*p* < 0.01, ratio paired two-tailed student’s *t*-test), confirming this notion.

How might stabilization of the A-loop in the upward configuration couple to channel gating? In this loop configuration, Lys671 is stabilized (and thus becomes well resolved) through its interaction with backbone carbonyls (Fig. 3A and B; and fig. S7) at the C-terminus of the TRP domain, a conserved α-helical motif that lies parallel to the inner membrane leaflet and has been implicated in gating of numerous TRP channel subtypes^16,17,21,25-27^. The positively charged side chain of Lys671 enhances the dipole moment of the TRP helix (fig. S7A and B), strengthening the positive electrostatic surface potential at its N-terminus (fig. S7C and D). We speculate that this promotes repulsion between TRP domains from neighboring subunits, thereby biasing the conformational equilibrium of the channel towards the open state by driving expansion of overlying S6 domains that constitute the lower gate (fig. S7E).

### Structurally conserved calcium control site

When TRPA1 was solubilized in LMNG, a robust density was seen at the bottom of each of the S2-S3 α-helices facing the cytoplasm (Fig. 4A and fig. S8A), where it is surrounded by residues Glu788, Gln791, Asn805, and Glu808, typical of calcium coordination sites^28^. Indeed, this configuration is strikingly similar to a calcium binding site seen in TRPM4 and TRPM8, with all relevant residues being conserved (Fig. 4A and B)^15,29^. Because channel purification was carried out in nominally calcium-free buffers, this density likely represents cellular calcium bound to a high affinity site.

**Fig. 4.**
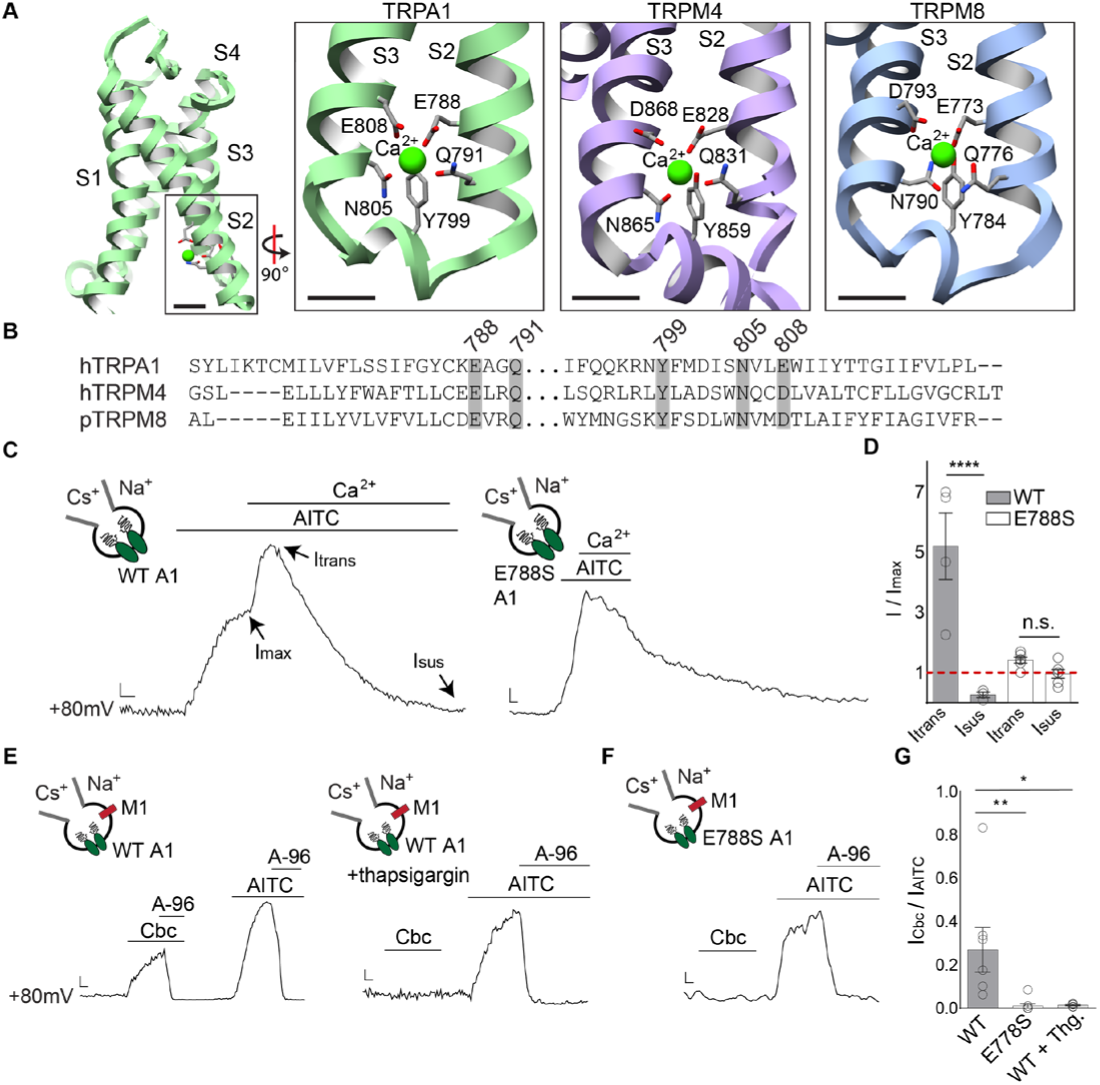
One site subserves three distinct calcium-dependent functions. **(A)** A calcium ion binds at the cytoplasmic end of S2-S3 transmembrane α-helices in the voltage sensor-like domain, analogous to calcium binding sites in TRPM4 and TRPM8. Scale bars = 5Å. **(B)** Sequence alignment between TRPA1, TRPM4 and TRPM8 reveals conservation of amino acid residues that coordinate calcium. **(C)** Normalized whole-cell recordings of TRPA1 WT and Glu788 Ser mutant currents following bath application of AITC (50 μM) and Ca^2+^ (2 mM). Data were extracted from 500msec. voltage ramps (−80 to 100mV) (n = 4-6 independent experiments; scale bars: x = 1 sec., y = arbitrary units (left) or x = 1 sec., y = arbitrary units (right). **(D)** Quantification of Ca^2+^ potentiation (I-transient, I_trans_, measured 2 sec. after Ca^2+^ application) and desensitization (I-sustained, I_sus_., measured 7.5 sec. post-Ca^2+^ application). ****, *p* < 0.0001 one-way ANOVA with post-hoc Holm-Sidak correction for multiple comparisons; n = 4-6 independent experiments; data displayed as mean ± S.E.M. **(E, F)** Normalized whole cell recordings for WT and Glu788Ser mutant TRPA1 channels co-expressed with M1 muscarinic acetylcholine receptor in response to bath applied carbachol (Cbc 100 μM), AITC (50 μM), and A-96 10 μM) with or without Thapsigargin (Thg,1 μM) pre-treatment (n = 5-9 independent experiments). **(G)** Quantification of responses (*, *p* < .05; ** *p* < 0.01, one-way ANOVA with post-hoc Holm-Sidak correction for multiple comparisons; n = 5-9 independent experiments). Data are displayed as the mean ± S.E.M.

Permeating calcium has two sequential effects on TRPA1, first enhancing currents and then promoting desensitization^10,19^. To determine if the calcium-binding site described above accounts for one or both of these effects, we expressed a TRPA1 mutant lacking one of the presumptive calcium coordination residues (Glu788Ser) and evaluated its response to physiological calcium. This mutant exhibited neither potentiation (in both whole-cell and excised-patch recordings) nor desensitization of AITC-evoked currents (observed with wild type channels (Fig. 4C and D; and fig. S8B), demonstrating necessity of this single site for these aspects of direct calcium modulation.

Like the canonical fly TRP channel, TRPA1 also functions as a receptor-operated channel that is activated downstream of metabotropic receptors that couple to phospholipase C signaling systems^5,6,9,10,30^. This mechanism of TRPA1 activation is likely mediated through consequent release of calcium from intracellular stores^10,19,31-35^ (although metabolism of phosphoinositide lipids has also been implicated^32-35^). To ask if this action converges on the same calcium site identified here, we co-expressed the M1 muscarinic receptor and TRPA1 in transfected HEK cells and recorded carbachol (an M1 agonist)-evoked TRPA1 responses in the absence of extracellular calcium. As previously shown^10^, carbachol elicited robust currents that were blocked by A-96 and not observed when intracellular calcium was depleted by pre-treatment with thapsigargin (Fig. 4E and G), reaffirming calcium as the critical second messenger. Most interestingly, the Glu788Ser mutant showed no response to carbachol (Fig. 4F and G), demonstrating that major regulatory actions of calcium on TRPA1 converge on this single calcium binding site.

## Discussion

Detectors of noxious stimuli must be tuned to function as early warning systems that recognize potentially injurious events before they elicit wholesale tissue damage. To accomplish this, they need to balance low threshold sensitivity with high fidelity. The two-step mechanism that we propose for electrophile-mediated TRPA1 activation (Fig. 5A) satisfies this requirement: Cys621 constitutes a high sensitivity site for initial irritant detection, which then initiates reorientation of the A-loop to prime a second site (Cys665) for reactivity and full channel activation. This two-step mechanism may be especially important when it comes to detecting and avoiding small volatile environmental toxicants such as acrolein or formalin (Fig. 5A). But as noted above, TRPA1 is unique in having evolved to recognize structurally diverse electrophiles, including larger endogenous agents such as reactive prostaglandins. In this case, low threshold and signal-to-noise considerations may be less relevant since rapid recognition and escape responses are not germane, and thus a single modification by bulkier electrophilic agonists may suffice (Fig. 5B).

**Fig. 5.**
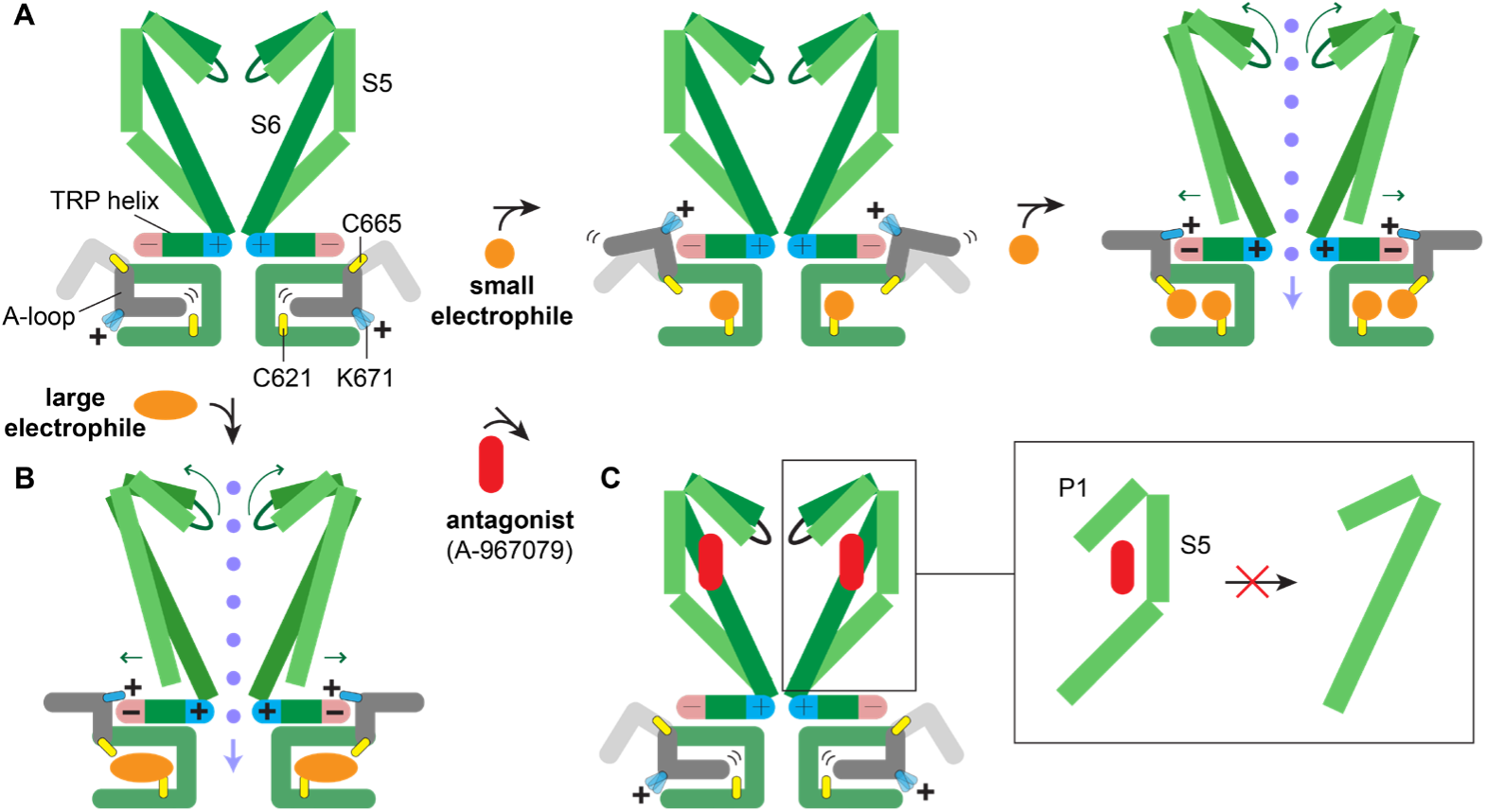
Model of TRPA1 activation. **(A)** Attachment of a small electrophile to Cys621 results in rearrangement of the A-loop to the ‘up’ position, bringing Cys665 into the reactive pocket. Modification of Cys665 by a second small electrophile stabilizes the A-loop and positions Lys671 at the C-terminus of the TRP helix. This increases positive electrostatic potential and charge repulsion at the N-termini of adjacent TRP helices, initiating conformational changes associated with dilation of the lower gate. These movements are coupled to widening of the upper gate and selectivity filter through straightening of the S5 helix. **(B)** Attachment of a large electrophile to Cys621 is sufficient to stabilize the A-loop in the ‘up’ conformation and activate the channel. **(C)** A-96 binds to the bent elbow region of S5, inhibiting the straightening of the α-helix required for channel gating.

In their roles as polymodal signal detectors, TRP channels are now recognized to contain two highly dynamic constrictions along their ion permeation pathway, one corresponding to a canonical lower gate and another at the level of the selectivity filter, which must be functionally coupled to mediate channel activation by diverse stimuli. In the case of TRPV1, vanilloid agonists likely enhance opening of both lower and upper gates, and we have proposed that this involves coupling of movement between pore helices in the selectivity filter and S5 transmembrane domains that ultimately shift positions of the pore lining S6 helices in the lower gate. However, we could only capture a fully opened TRPV1 structure in the presence of both a vanilloid agonist that opens the lower gate and a spider toxin that stabilizes the upper gate. Thus, while we surmise from functional studies that allosteric coupling connects these two restrictions, our TRPV1 structures did not provide direct evidence for this effect. In the case of TRPA1 we see a fully open state in the presence of agonists that target a region below the transmembrane core, providing more direct support for an allosteric coupling mechanism. More generally, our findings suggest that the upper restriction in the voltage-gated ion channel superfamily has evolved to serve a range of physiologic functions: in voltage-gated potassium channels and some TRP channels (*i.e.*, TRPM2) this region is devoted largely to ion selection^26,36^, whereas in TRPV1 it serves as a regulated gate^16-18^. In the case of TRPA1, our results suggest an intermediate function of a regulated region controlling dynamic ion selectivity, which likely underlies the differential ability of activators to elicit pain and neurogenic inflammation^21^.

Another common theme among TRP channels is their modulation by calcium, which serves as a sensitizing and/or desensitizing second messenger. Modulation may be mediated through a calcium binding partner, such as calmodulin in the case of TRPV5 or TRPV6^37-42^, or through direct interaction with the channel. The calcium binding site that we identify in TRPA1 supports an emerging picture of a conserved structural motif initially seen in TRPM channel subtypes (Fig. 4A and B)^15,29,43-45^. In all cases, this is a key site for detecting elevated intracellular calcium resulting from channel activation and/or store release. For TRPA1 this accounts for multiple calcium-dependent processes, including sensitization, desensitization, and coupling to metabotropic receptors. For TRPM8, this conserved calcium binding pocket is adjacent to a ligand binding site and calcium is required for channel activation by the synthetic super-cooling agent, icilin^15,43^. Whether calcium similarly regulates activation of TRPA1 by as-yet undiscovered ligands or post-translational modifications^46^, or how calcium binding is translated to gating movements, remain intriguing questions for future studies.

## Materials and Methods

### Protein expression and purification

Human TRPA1 was expressed and purified as described previously with slight modifications^11^. Briefly, N-terminal-tagged MBP-TRPA1 under the control of a CMV promoter was expressed in HEK293F cells using the BacMam system (Thermo Fisher). Protein expression was induced at cell density 1.5-2.5 × 10^6^ cells ml^-^ with media supplemented with 5 mM sodium butyrate and 5 µM ruthenium red. After 16 hr. at 37 °C, cells were harvested by centrifugation at 1,000 × *g* for 10 mins. Cells were resuspended in lysis buffer (50 mM HEPES, pH 8.0, 150 mM NaCl, 1 mM IP_6_, 1 mM DTT, 1 mM EDTA, protease inhibitors) and lysed by sonication. Cell membranes were isolated by ultracentrifugation at ∼180,000 × *g* for 45 mins and the pellet was resuspended in lysis buffer before multiple passes through a glass homogenizer. Cell membranes were solubilized by addition of LMNG or Cymal5-NG (Anatrace) to a final concentration of 0.5 % (w/v) and rocking at 4°C for 1 hr. Insoluble debris was removed by centrifugation at ∼35,000 × *g* for 20 mins. and the supernatant passed over amylose beads by gravity flow at 4°C. The beads were washed with wash buffer (50 mM HEPES, pH 8.0, 150 mM NaCl, 1 mM IP_6_, 1 mM DTT, 0.005% LMNG/Cymal5-NG) and eluted with wash buffer supplemented with 10 mM maltose.

For exchange into amphipol, PMAL-C8 was added to the detergent-solubilized sample in a 3:1 PMAL:protein ratio. The solution was mixed by rocking at 4°C for 1 hr. before addition of 100 μl bio-beads (Biorad) ml^-^ of protein sample. The sample with bio-beads was mixed by rocking at 4°C overnight. The bio-beads were spun down in a tabletop centrifuge at 100 × *g* and the supernatant analyzed by size exclusion chromatography in detergent-free buffer (50 mM HEPES, pH 8.0, 150 mM NaCl, 1 mM IP_6_, 1 mM DTT). Peak fractions corresponding to TRPA1 were pooled and concentrated 0.5 - 2 mg ml^-^ in a 100K MWCO centrifuge concentrator.

### Agonists and antagonists

To acquire samples of activated TRPA1 for structural studies, cell membranes containing TRPA1 was incubated with 100 µM iodoacetamide (IA) or BODIPY-IA(Millipore Sigma) for 10 min. prior to protein purification. Samples of antagonist-bound TRPA1 were acquired by incubating the purified channel with 10 µM A-967079 (A-96) (Tocris) for 10 mins. prior to cryo-EM grid preparation.

### Rationale and interpretation of solubilization conditions

TRPA1 protein was imaged in both PMAL-C8 amphipol and LMNG detergent. When apo TRPA1 was solubilized in PMAL-C8, we observed two conformational states (closed and open) within the same sample (Fig. 1A), both of which lacked ions in the calcium binding site. However, the A-loop could not be modelled, consistent with the dynamic nature of the loop and the ability of the channel to adopt different states (fig. S3). Treatment with IA stabilized the A-loop in the ‘up’ conformation in conjunction with an open pore, which we therefore designated an activated state of the channel. Addition of agonist before detergent extraction from membranes was crucial for isolating the channel in the activated state, presumably because solubilization in detergent or amphipol prevents dynamic exchange between different channel conformations. Consistent with this notion, when A-96 was added after channel solubilization in PMAL-C8, the antagonist recognized, but did not shift the equilibrium towards the closed conformation.

In LMNG, the transmembrane core of the channel adopted a closed conformation under both apo and BIA conditions, possibly due to locking of the transmembrane domain by LMNG and/or desensitization of the channel by bound ions in the calcium binding site. Consistent with a strongly-biased closed state, the A-loop was clearly observed in the ‘down’ (closed) conformation of the apo channel. However, modification of TRPA1 by BIA stabilized the A-loop in the ‘up’ (activated) conformation in conjunction with a closed pore, which likely represents an intermediate state of the channel following electrophile modification.

### Cryo-EM data collection and processing

Samples for cryo-EM were prepared by applying 4 µl of purified TRPA1 to 1.2/1.3 holey carbon grids (Quantifoil) and blotting for 8-12 sec. in a Vitrobot Mark IV (Thermo Fisher) prior to plunge freezing in liquid ethane. For multi-shot imaging, samples were prepared on 2/2 holey carbon grids (Quantifoil) and blotted for 6-8 s. Cryo-EM samples were imaged on Polara and Titan Krios microscopes (Thermo Fisher, see Supplementary Figure 1d for details). Movies were drift-corrected using *MotionCor2*^48^ and CTF parameters estimated with *gctf* ^49^. Particle images were selected from micrographs using *gautomatch* (MRC-LMB, https://www.mrc-lmb.cam.ac.uk/kzhang/) and extracted in *Relion*^50^. 2- and 3-D classification of particle images was performed in *cryoSPARC*^51^ and 3D maps refined in *cryoSPARC* and *cisTEM*^52^. Conversion of data from *cryoSPARC* to *Relion* and generation of orientation distribution plots were performed using *pyem*^*53*^. Directional Fourier shell correlations of cryo-EM maps were performed as previously described^54^.

### Model building and analysis

Cryo-EM maps were visualized with UCSF Chimera^55^. Atomic models were built into the cryo-EM maps with Coot^56^ using the previously published structure of TRPA1 as a starting model (PDB-ID: 3J9P). The models were refined over multiple rounds using PHENIX Real Space Refinement^57^. Ligands were built using Coot and their geometric restraints calculated with Phenix eLBOW^58^. Cysteine pK_a_s were calculated in H++^59^ and the resultant values used to solve the Henderson-Hasselbalch equation^60,61^ to determine the percentage free sulfhydryl (fig. S6J). Electrostatic surface potentials for the Apo-LMNG and BODIPY-IA-LMNG structures were calculated in APBS^47^ using an AMBER forcefield and are displayed at ± 10 kT/e^-^ (Fig. 2D; and fig. S5 and 8).

### Molecular Biology and Cell Culture

We subcloned full-length wild-type human TRPA1 into the mammalian/oocyte expression vector pMO, or pcDNA3.1^+^ containing an N-terminal eGFP tag, which served as templates for all physiology experiments^11,62,63^. Constructs generated from these templates were produced by Gibson assembly (New England Biolabs) and verified by sequencing.

HEK293T (ATCC) cells were cultured at 37 °C, 5% CO_2_ in DMEM Complete (DMEM-C; Dulbecco’s modified Eagle’s medium containing 10% (v/v) heat-inactivated calf serum, 100 U ml^-1^ Penicillin G and 0.1 mg ml^-1^ Streptomycin). For heterologous expression of ion channels, HEK293T cells were transfected with 0.25-1 μg of plasmid DNA combined with 3x (w/w) Lipofectamine 2000 (Thermo-Fisher) for 4-8 hrs. in Opti-MEM (Thermo Fisher) before being plated onto coverslips coated with 0.01% poly-L-Lysine (MW 70-150,000, MilliporeSigma). Cells were then adhered to these coverslips for at least 12 hr. before use in patch-clamp experiments.

### Electrophysiology

Capillary pipettes from borosilicate glass with filament (O.D × I.D, 1.10 × 0.86 mm, Sutter Instruments) were fashioned and fire-polished to a resistance of 5-10 mΩ for whole-cell and excised-patch patch-clamp recordings. Electrophysiological data were collected at RT using an Axopatch 200B amplifier (Axon Instruments) and digitized with a Digidata 1550B (Axon Instruments). Voltage protocols were delivered and resulting currents monitored on-line with pClamp10 (Molecular Devices), then analyzed off-line in pClamp or Prism (GraphPad). All electrophysiological recordings and pharmacological manipulations were carried out under laminar flow using a pressure-driven micro-perfusion system (SmartSquirt, Automate Scientific).

For whole-cell recordings, the bath solution consisted of Ca^2+^-free Ringer’s solution (140 mM NaCl, 10 mM HEPES, 5 mM KCl, 2 mM MgCl_2_, and 10 mM Glucose; pH 7.4 with NaOH; 290-300 mOsm kg^-1^data), data were digitized at 10 kHz and filtered at 1 kHz. The internal solution contained 140 mM CsMeSO_4_; 10 mM HEPES; 1 mM MgCl_2_; and, as indicated, 0 or 1 mM EGTA. The pH was set to 7.2 with CsOH and osmolarity to 300-310 mOsm kg^-1^ with sucrose. Unless otherwise stated in the figure legends, analysis of IA- and BIA-evoked TRPA1 currents in whole-cell patch clamp mode was carried out at V_hold_ = −80 mV. Current-voltage relationships were then measured at steady state in response to voltage steps (500 msec.) from −80 to 80 mV in 10 mV increments using on-line leak subtraction (P/4). Tail-currents were then evoked by a brief (250 msec.) test pulse of −120 mV. The decay-time constant τ for each tail-current was determined by fitting a one-phase exponential decay function in pClamp to the tail-current obtained following the 80 mV step^13^. For analysis of Ca^2+^-modulation, TRPA1 currents were continuously monitored over a 500 msec. voltage ramp from −80 to 100 mV.

Inside-out excised-patch recordings were carried out in symmetrical solutions of 150 mM NaCl, 10 mM HEPES, 2 mM EGTA, 1 mM MgCl_2_; pH 7.3 with NaOH; 300-310 mOsm kg^-1^ at a constant holding voltage of −40 or −60 mV (Fig. S9B and C) (fig. S9D); sampled at 20 kHz; and filtered at 2 kHz. For measuring monovalent cation conductance, the solutions were as above, minus Mg^2+^. For divalent cation conductance measurements, the bath solution contained: 75 mM CaCl_2_, 10 mM HEPES, pH 7.3 with Trizma; 300-310 mOsm kg^-1^. The pipette solution contained 150 mM NaCl, 10 mM HEPES, 2 mM EGTA; pH 7.3 with NaOH; 300-310 mOsm kg^-1^. The reversal potential was determined as the minimum of the standard deviation of the average trace resultant from at least 10 sweeps of a 500 msec. ramp from −100 to 80 mV under a given pharmacological condition. Relative permeability ratios calculated using this value were then used to solve the Goldman-Hodgkin-Katz equation^13,21,22^.

### Statistics and Experimental Design

Herein we summarized electrophysiological data using the mean ± SEM unless otherwise noted. We carried out statistical testing in Prism (GraphPad) using either a ratio paired two-tailed student’s *t* test, or One-way ANOVA with a *post-hoc* Holm-Sidak correction for multiple comparisons, as indicated in the Figure legends. *A priori*, we set α = 0.05 and represent statistical significance as: * *P* < 0.05, ** *P* < 0.01, and **** *P* < 0.0001. Parametric significance tests assuming equal variance and a normal distribution of data means are justified given the experimental design; they are standard tests for similar experiments. We selected sample sizes for all experiments based on our laboratory and others’ experience with similar assays, and in consideration of reagent availability and technical feasibility. We made no predetermination of sample size and thus carried out a minimum of independent experiments required for statistical significance and reproducibility.

## End Matter

### Author Contributions and Notes

J.Z. designed and executed biochemical and cryo-EM experiments, with early collaborative contribution and guidance on TRPA1 expression and purification from C.E.P. J.V.L.K. designed and carried out physiology experiments. J.Z., J.V.L.K, Y.C., and D.J. conceived the project, interpreted the results, and wrote the manuscript. The authors declare no competing interests.

## Acknowledgments

We thank all members of the Julius and Cheng labs for thought-provoking discussion and critical reading of the manuscript. Some data for this study were collected at the Toronto High-Resolution High-Throughput cryo-EM facility, supported by the Canada Foundation for Innovation and Ontario Research Fund. This work was supported by an American Heart Association Postdoctoral Fellowship (J.Z.), a Banting Postdoctoral Fellowship from the Canadian Institutes of Health Research (J.Z.), an NSF Graduate Research Fellowship (No. 1650113 to J.V.L.K), a UCSF Chuan-Lyu Discovery Fellowship (J.V.L.K), a Helen Hay Whitney Foundation Postdoctoral Fellowship (C.E.P.) and grants from the NIH (R35 NS105038 to D.J; R01 GM098672, S10 OD021741, and S10 OD020054 to Y.C.; T32 HL007731 to C.E.P.; and T32 GM007449 to J.V.L.K). Y.C. is an investigator of the Howard Hughes Medical Institute.

---

**Fig. S1.**
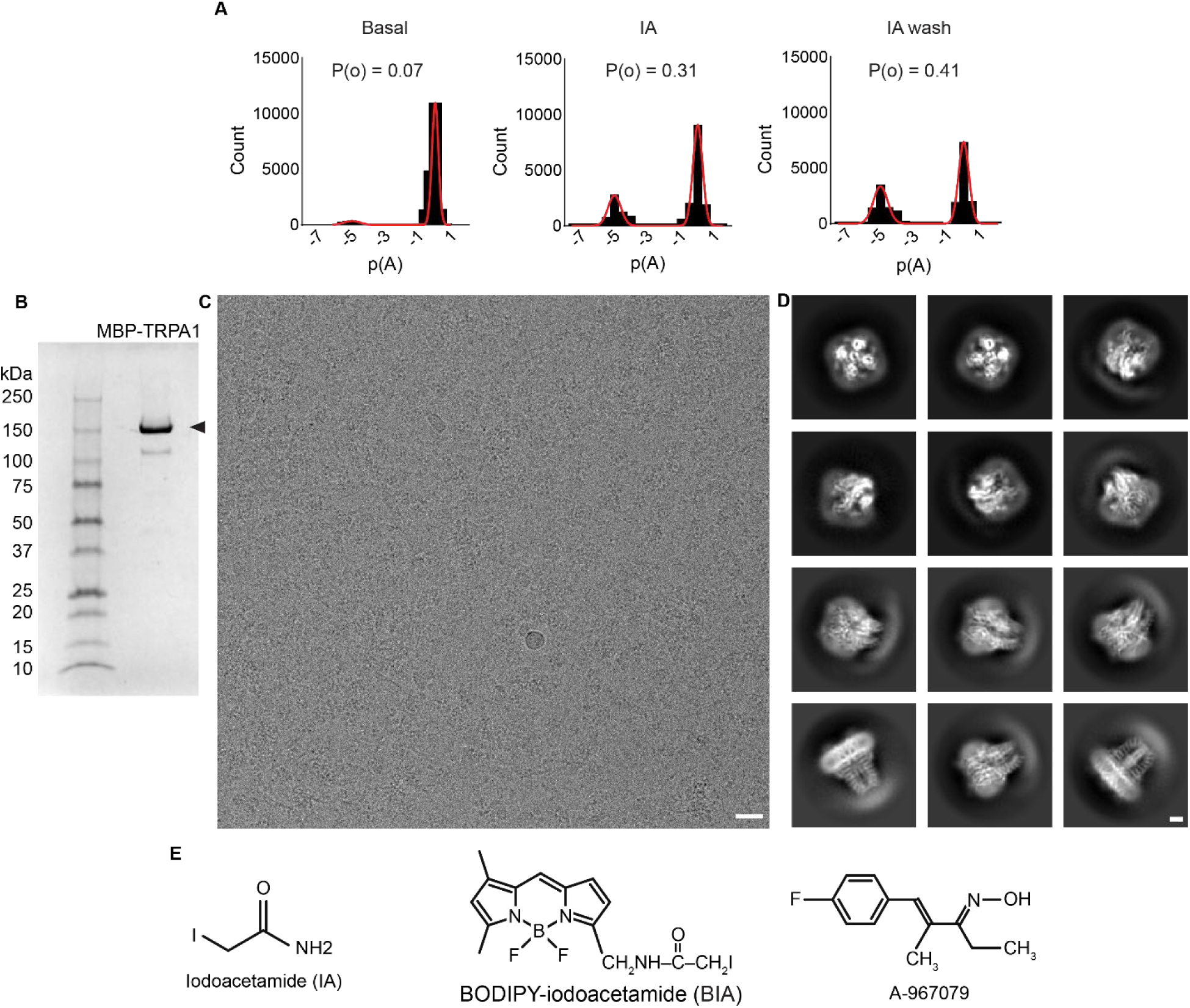
Pharmacology and Cryo-EM Data Collection and Processing of TRPA1. **(A)** All points histograms depicting the change in open probability (P(o)) in a single TRPA1 channel in response to IA-application. Data represent n = 9 independent excised inside-out patches. V_hold_ = −40 mV **(B)** SDS-PAGE showing MBP-TRPA1 (arrowhead) after pull-down and elut5ion from amylose beads. **(C)** Cryo-EM image of MBP-TRPA1. Scale bar: 20 nm. **(D)** 2D classification of cryo-EM particle images showing TRPA1 in different orientations. Scale bar: 25 Å. **(E)** Cryo-EM data collection parameters. **(F)** Pharmacological agents used in this study.

**Fig. S2.**
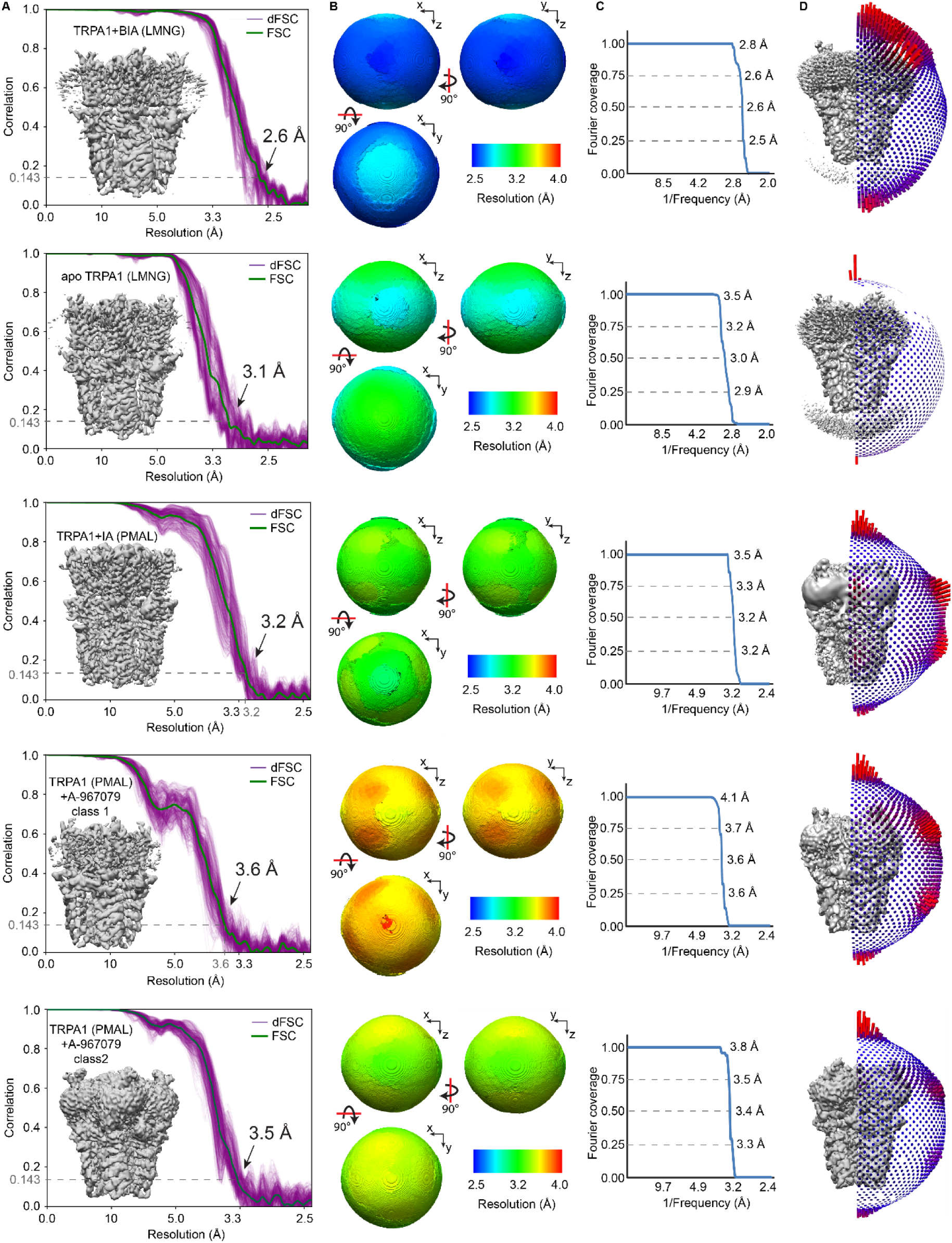
Fourier shell correlation of cryo-EM maps and orientation distribution of particle image views. **(A)** Fourier shell correlation and 1D directional Fourier shell correlation plots. **(B)** 3D representations of the directional Fourier shell correlation. **(C)** Fourier space covered, based on dFSC at 0.143. **(D)** Orientation distribution of particle image refinement angles.

**Fig. S3.**
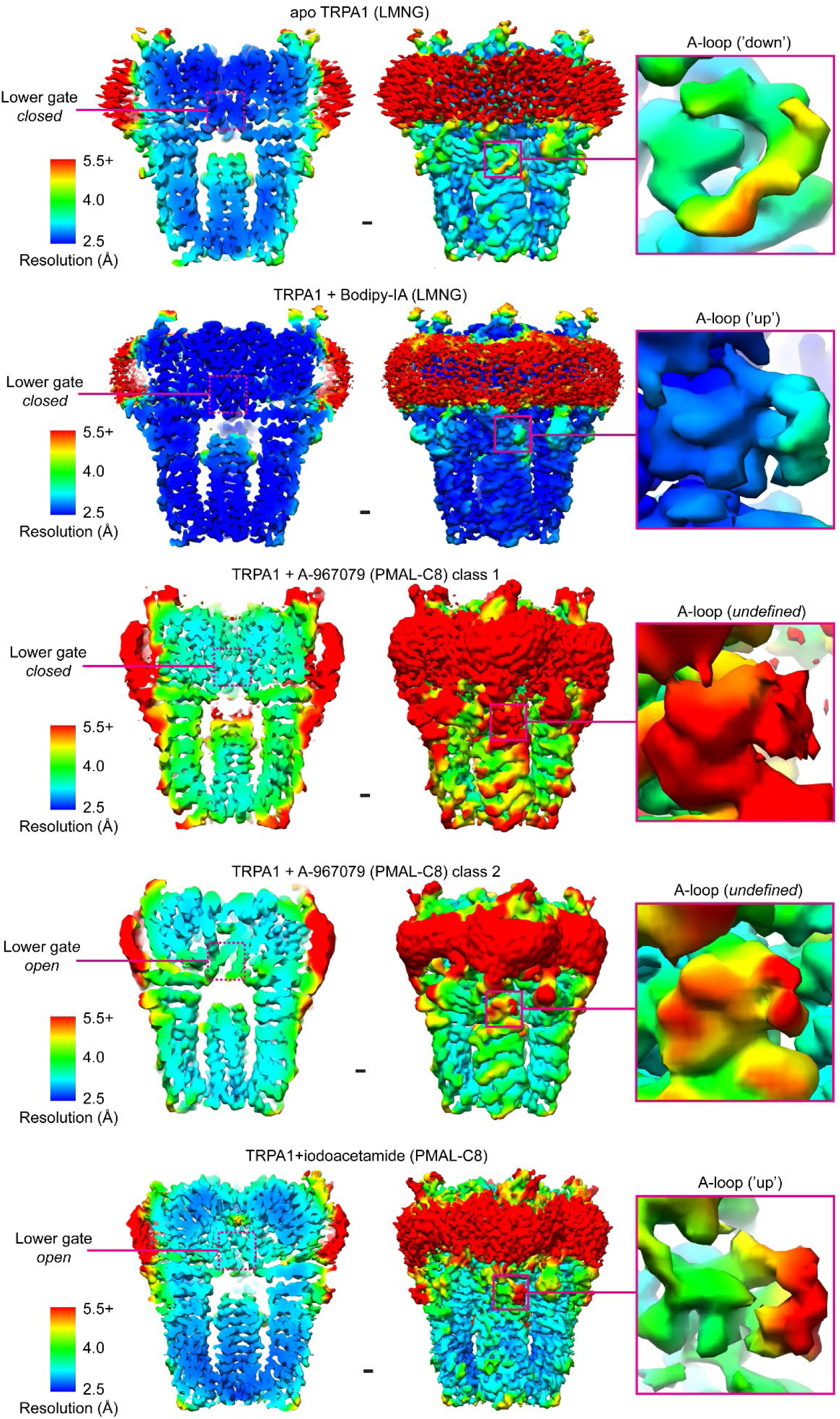
Local resolution of TRPA1 cryo-EM maps. The A-loop is lower resolution compared to surrounding map regions, indicating its dynamic nature. In the activated conformation (TRPA1+iodoacetamide), the bottom of S6 is lower resolution compared to surrounding regions, indicating structural flexibility at the level of the lower gate. Scale bars: 5 Å.

**Fig. S4.**
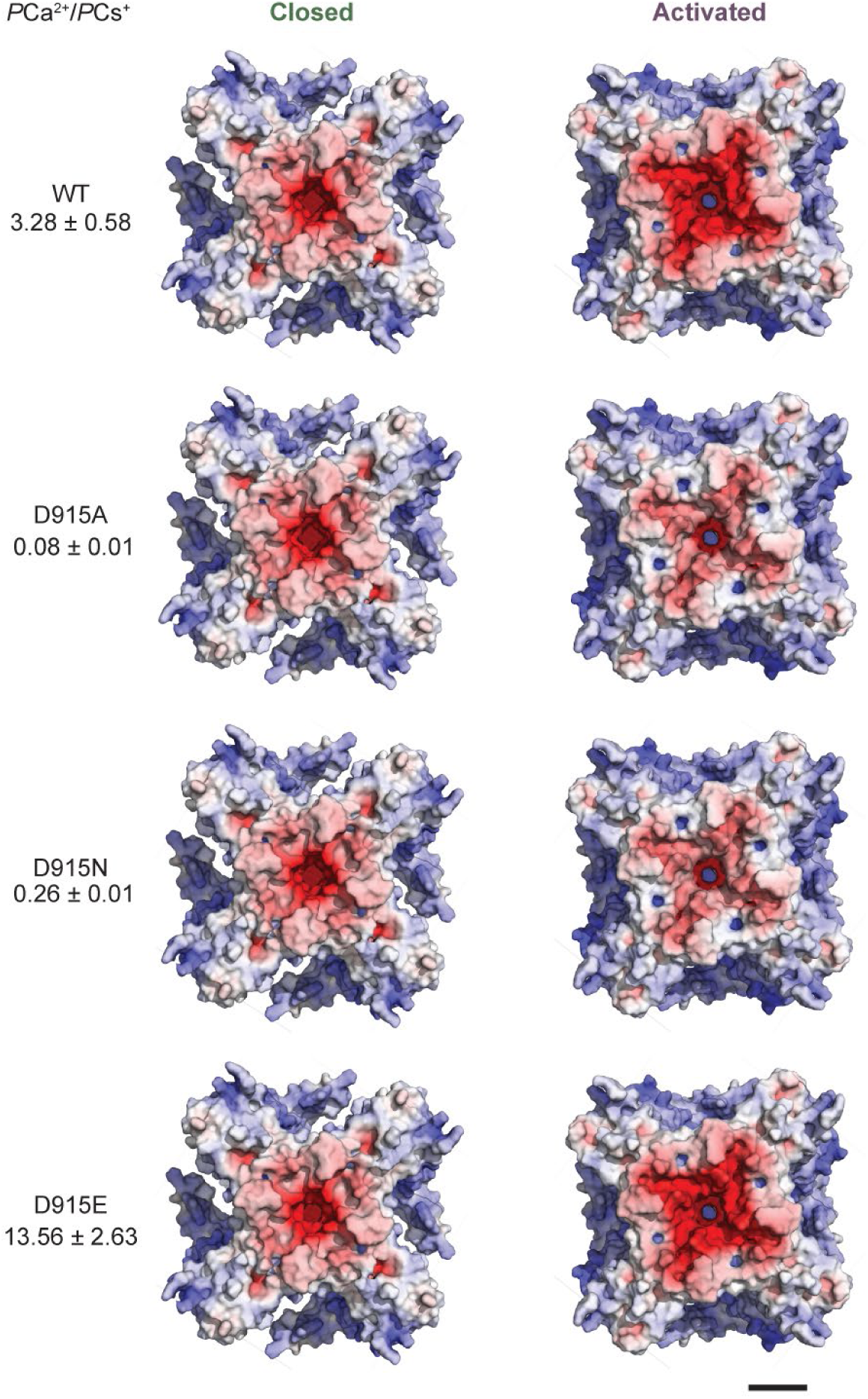
Surface charge distribution of TRPA1’s extracellular face. Electrostatic potential maps were calculated in APBS and are displayed at ± 10 kT/e^-^. *In silico* mutations of Asp915 were modeled and experimentally determined relative permeability ratios for these mutations sourced from Wang *et al.* (ref. 19). Scale bar: 30 Å

**Fig. S5.**
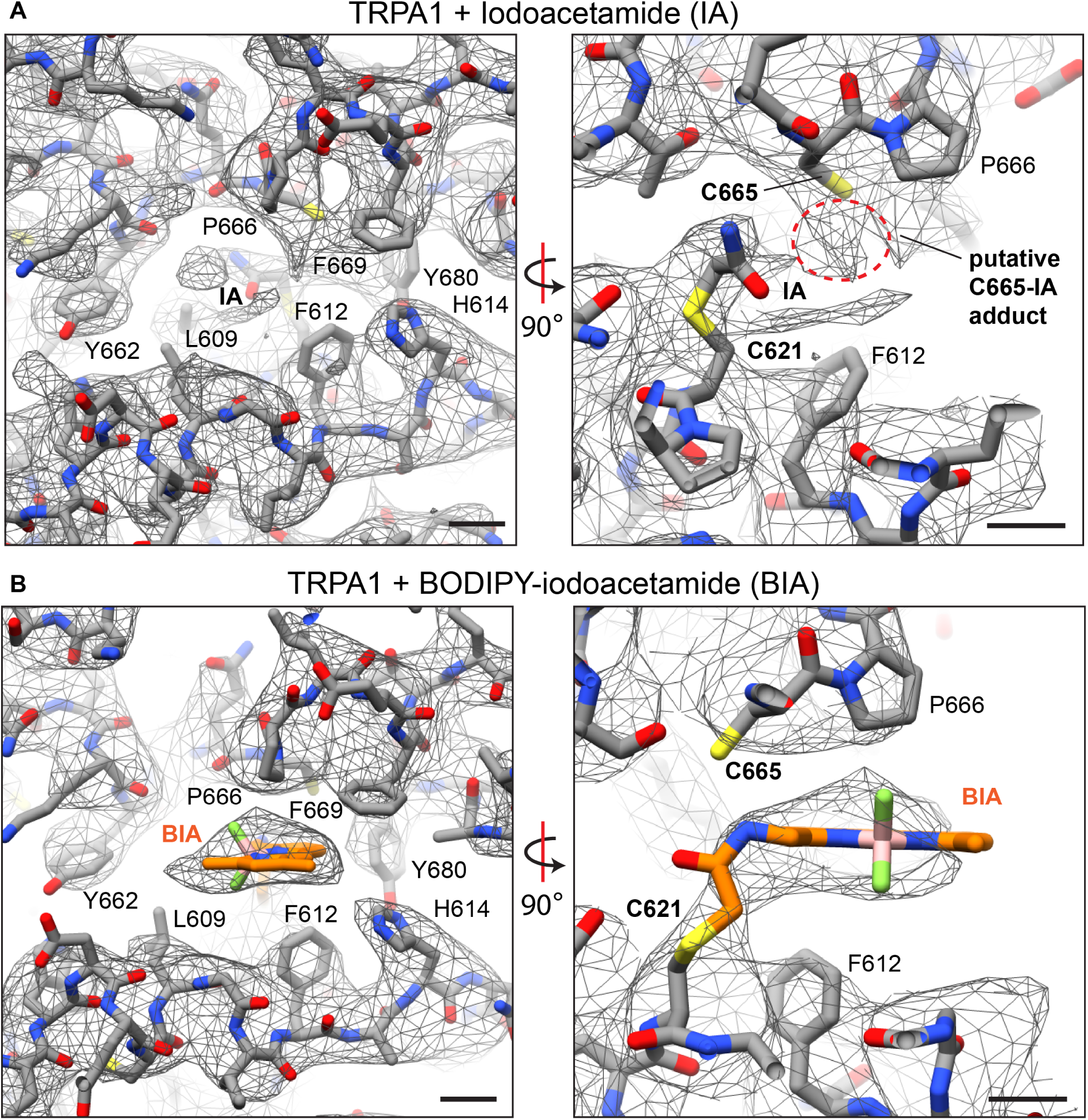
Map density of agonists. (A) Strong density is observed for iodoacetamide bound to Cys621. Weaker density is observed next to Cys665, which indicate that some of the channels may be modified by agonist at this site. (B) Clear density for BODIPY-iodoacetamide (BIA) is observed bound to Cys621. No additional density is observed next to Cys665 in this case.

**Fig. S6.**
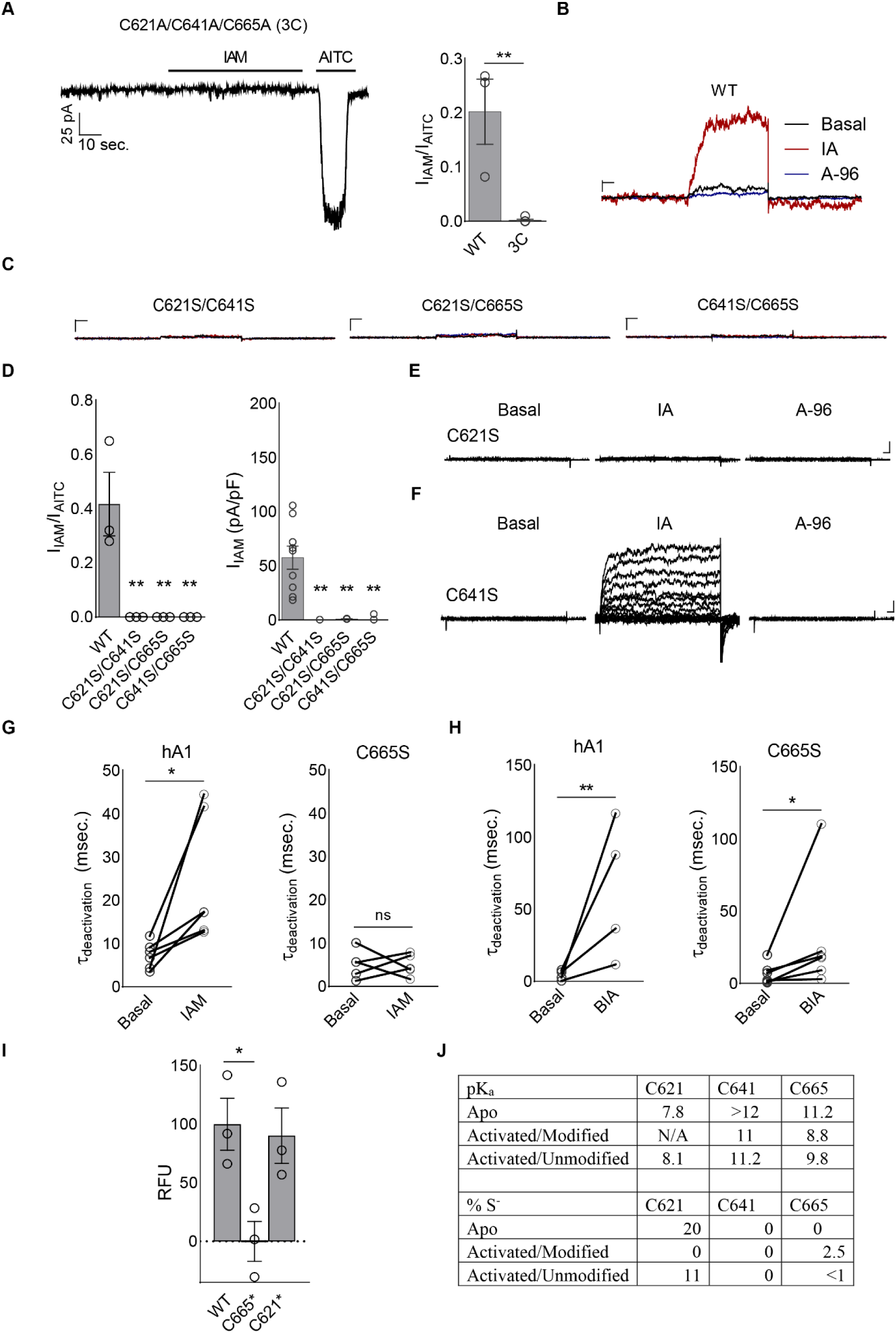
Electrophysiological Characterization of TRPA1 activation by IA and BIA. **(A)** IA (100μM) activates TRPA1 through covalent modification of cysteines; AITC (250μM). Data represent 3-9 independent experiments. Statistical significance is reported as the result of an unpaired two-tailed student’s *t*-test. V_hold_ = −80 mV**. (B-D)** No single cysteine is sufficient for TRPA1 activation by IA. Data represent 3-9 independent experiments. Data were acquired in whole-cell patch clamp mode and reflect the results of 500msec. test pulse (80mV). V_hold_ = −80 mV. Doses: IA (100 μM), A-967079 (10 μM). Scale bars: x, 50 msec.; y, 100 pA. (**E)** Cys621Ser display complete loss of IA sensitivity. Data represent n = 5 independent experiments and **(F)** Cys641Ser retains full sensitivity to IA. Data represent n = 5 independent experiments. Data were acquired in whole-cell patch clamp mode and reflect the results of 500 msec. test pulses from −80 to 80mV. V_hold_ = −80mV. Doses: IA (100μM), A-96 (10μM). Scale bars: x, 25 msec.; y, 100 pA. **(G)** Quantification (n = 4 - 6 cells) of changes in IA and **h**, BIA-evoked TRPA1 tail-current decay time constants in WT and Cys665Ser TRPA1. Data were acquired in whole-cell patch clamp mode after a 500msec. pre-pulse (80 mV) followed by a 250msec. test pulse (−120mV). V_hold_ = −80 mV. Statistical significance is represented as the results of a ratio paired two-tailed student’s t-test. **(I)** Binding of BIA to TRPA1 Cys64 Ser/Cys665Ser double mutant is similar to wildtype. Statistical significance is represented as the results of one-way ANOVA with post-hoc Holm-Sidak correction for multiple comparisons. **(J)** TRPA1 cysteine pK_a_ values and deduced proportion of thiolate in the Apo state (PDB-ID: 6V9W), and IA-bound (‘Activated,’ PDB-ID: 6V9V) state in the presence or absence of covalent modification at Cys621.

**Fig. S7.**
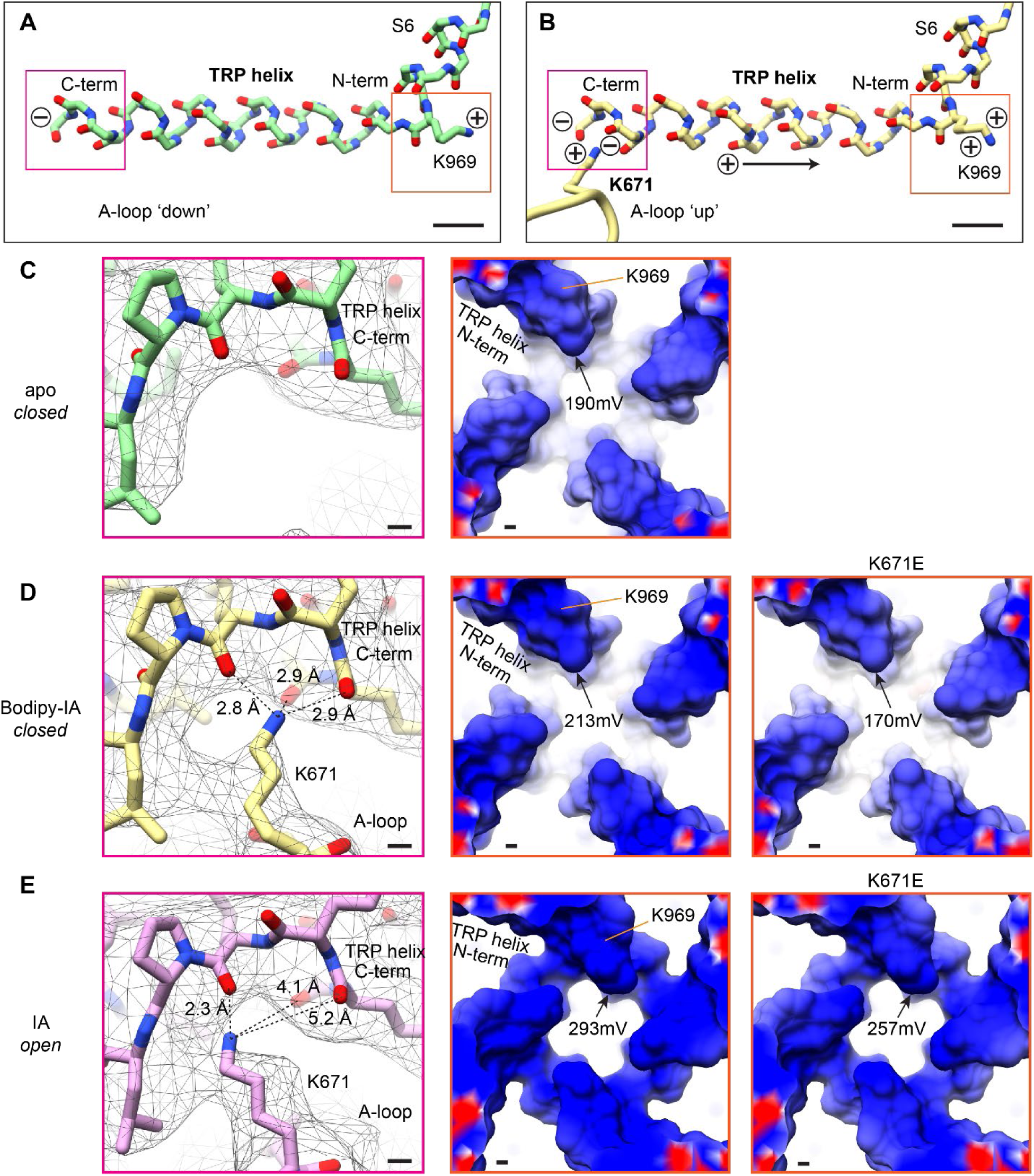
Positive electrostatic potential below the lower gate. **(A)** The TRP helix forms an electric dipole with electro-positive Lys969 at the N terminus and electro-negative carbonyl oxygens at the C terminus. **(B)** When the A-loop is oriented in the ‘up’ position, Lys671 is coordinated by the carbonyl oxygens at the C terminus of the TRP helix and increases its dipole moment to enhance the positive electrostatic potential at the N terminus. **(C)** The C-terminal carbonyl oxygens of the TRP helix form a pocket that is unoccupied in the apo channel. **(D)** Coordination of Lys671 with the carbonyl oxygens at the TRP helix C terminus increases the positive electrostatic potential at the TRP helix N terminus. *In silico* substitution of Lys671 with Glu decreases the electrostatic potential of the TRP helix. **(E)** Conformational changes associated with pore dilation further increase the positive electrostatic potential of the TRP domain.

**Fig. S8.**
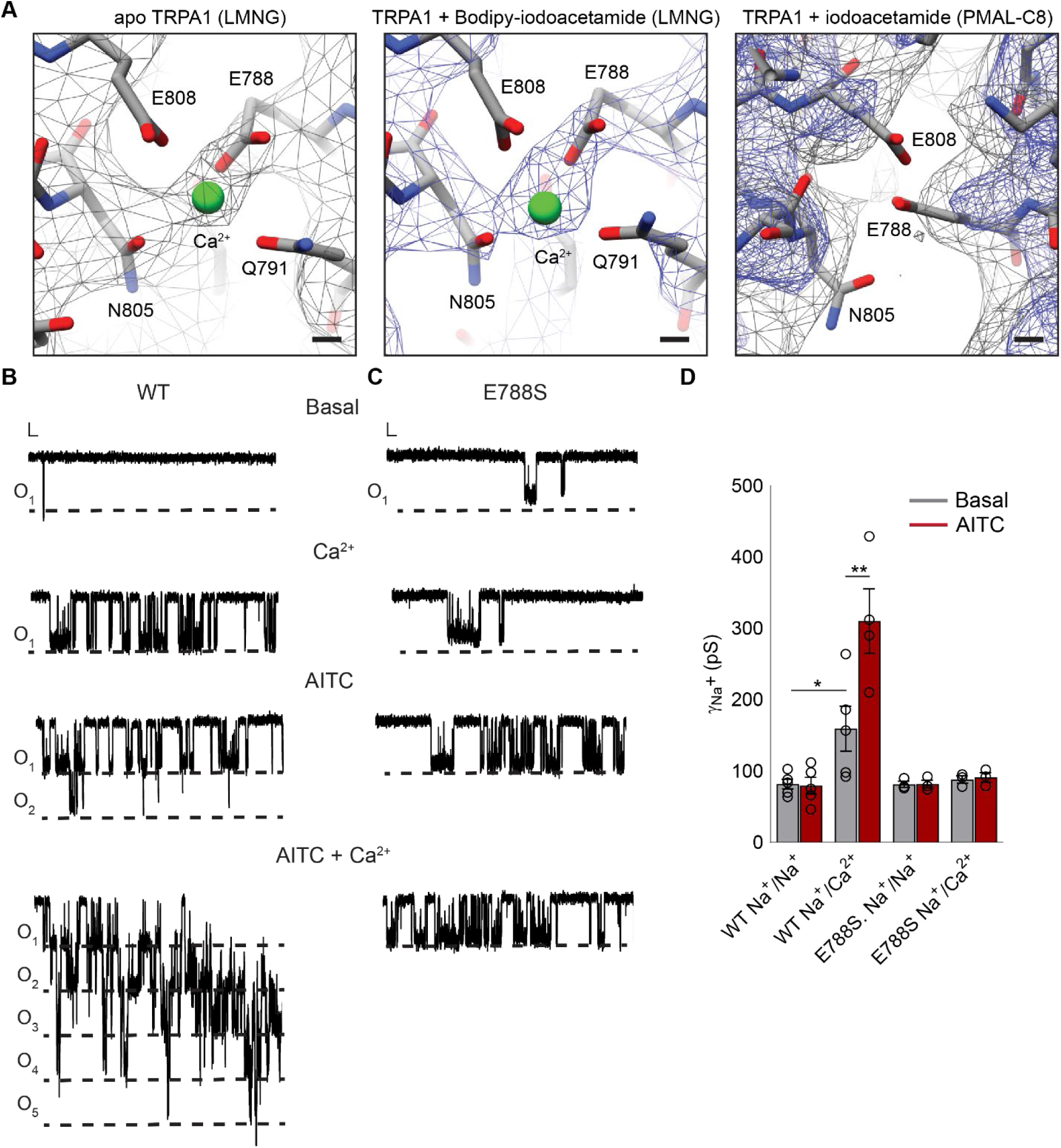
Calcium map densities and single-channel recordings of Ca^2+^ modulation. **(A)** Calcium is bound in both apo (σ = 4) and agonist-treated (σ = 8) samples in LMNG detergent, with Glu788 and Asn805 displaying the most robust densities coordinating calcium. No density for calcium is observed for the channel in amphipol (gray, σ = 4; blue, σ = 8). **(B-C)** Single channel recording from an excised inside-out patch of TRPA1 activation and potentiation by Ca^2+^. Data represent n = 3 independent experiments. Scale bar: x, 10 msec.; y, 2 pA. V_hold_ = −60mV. Doses: Ca^2+^, 2mM; AITC, 50μM. **(D)** Excised inside-out patch conductance measurements of TRPA1 in the presence or absence of intracellular Ca^2+^ in the basal and AITC (50μM)-activated states. Data are represented as the mean ± S.E.M and statistical significance represents the results of a one-way ANOVA with post-hoc Holm-Sidak correction for multiple comparisons, n = 3 −5 independent experiments.

**Fig. S9.**
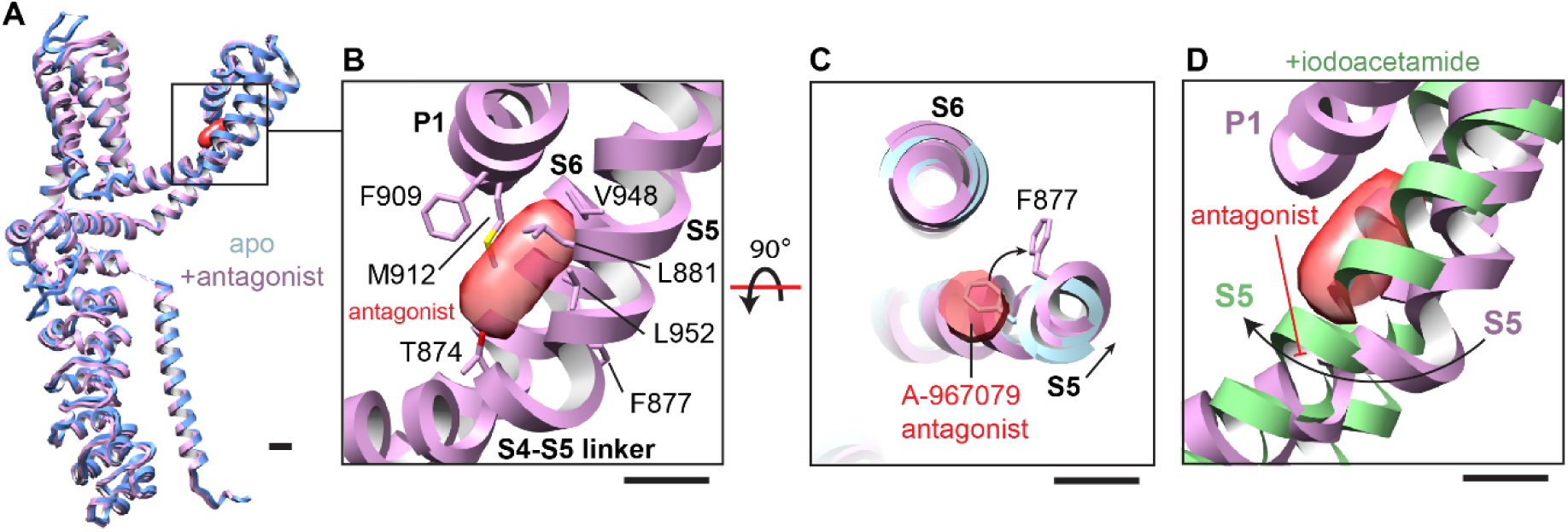
Binding of A-967079 to TRPA1. **(A)** The overall architecture of apo and antagonist-bound TRPA1 is similar, representing a closed state. **(B)** A-967079 binds at the elbow of S5, sandwiched between S6 and P1. **(C)** Binding of A-967079 results in a slight shift in S5 and repositioning of Phe877. **(D)** The antagonist is in an ideal position to block the straightening of the S5 elbow and inhibit channel gating.

**Table S1.** Cryo-EM data collection, refinement and validation statistics.

|  | TRPA1+BIA<br>(LMNG)<br>(EMD-21127)<br>(PDB 6V9V) | TRPA1 apo<br>(LMNG)<br>(EMD-21128)<br>(PDB 6V9W) | TRPA1+IA<br>(PMAL-C8)<br>(EMD-21129)<br>(PDB 6V9X) | TRPA1<br>+A-967079<br>(PMAL-C8)<br>Class 1<br>(EMD-21130)<br>(PDB 6V9Y) | TRPA1<br>+A-967079<br>(PMAL-C8)<br>Class 2<br>(EMD-21131) |
| --- | --- | --- | --- | --- | --- |
| **Data collection and processing** |  |  |  |  |  |
| Magnification | 22,500 | 75,000 | 31,000 | 31,000 | 31,000 |
| Voltage (kV) | 300 | 300 | 300 | 300 | 300 |
| Electron exposure (e-/Å<sup>2</sup>) | 25 | 30 | 40 | 40 | 40 |
| Defocus range (µm) | 0.8-2.0 | 0.8-2.0 | 1.3-2.5 | 1.3-2.5 | 1.3-2.5 |
| Pixel size (Å) | 1.05 | 1.06 | 1.22 | 1.22 | 1.22 |
| Symmetry imposed | C4 | C4 | C4 | C4 | C4 |
| Initial particle images (no.) | 374,863 | 242,399 | 456,208 | 296,991 | 296,991 |
| Final particle images (no.) | 212,751 | 130,595 | 190,112 | 30,321 | 78,959 |
| Map resolution (Å) | 2.6 | 3.1 | 3.3 | 3.6 | 3.5 |
| FSC threshold | 0.143 | 0.143 | 0.143 | 0.143 | 0.143 |
| Map resolution range (Å) | 2.3-4.1 | 2.6-5.2 | 2.8-5.6 | 3.2-8.5 | 3.1-6.5 |
| **Refinement** |  |  |  |  |  |
| Initial model used (PDB code) | 3J9P | 3J9P | 3J9P | 3J9P |  |
| Model resolution (Å) | 2.8 | 3.4 | 3.4 | 3.8 |  |
| FSC threshold | 0.5 | 0.5 | 0.5 | 0.5 |  |
| Model resolution range (Å) | 357-2.8 | 360-3.4 | 311-3.4 | 311-3.8 |  |
| Map sharpening *B* factor (Å<sup>2</sup>) | -25 | -25 | -100 | -100 |  |
| Model composition |  |  |  |  |  |
| Non-hydrogen atoms | 18,976 | 18,812 | 18,084 | 17,716 |  |
| Protein residues | 2,384 | 2,388 | 2,280 | 2,248 |  |
| Ligands | 8 | 4 | 4 | 0 |  |
| *B* factors (Å<sup>2</sup>) |  |  |  |  |  |
| Protein | 89 | 165 | 55 | 112 |  |
| Ligand | 112 | 216 | 57 | 0 |  |
| R.m.s. deviations |  |  |  |  |  |
| Bond lengths (Å) | 0.008 | 0.008 | 0.004 | 0.009 |  |
| Bond angles (°) | 0.904 | 0.965 | 0.880 | 1.179 |  |
| Validation |  |  |  |  |  |
| MolProbity score | 1.57 | 1.62 | 1.48 | 1.99 |  |
| Clashscore | 3.47 | 4.53 | 3.52 | 6.53 |  |
| Poor rotamers (%) | 1.2 | 1.21 | 1.24 | 2.13 |  |
| Ramachandran plot |  |  |  |  |  |
| Favored (%) | 94.6 | 95.26 | 96.07 | 94.38 |  |
| Allowed (%) | 5.4 | 4.74 | 3.93 | 5.62 |  |
| Disallowed (%) | 0.0 | 0.0 | 0.0 | 0.0 |  |

